# Phylodynamic assessment of control measures for highly pathogenic avian influenza epidemics in France

**DOI:** 10.1101/2021.06.23.449570

**Authors:** Debapriyo Chakraborty, Claire Guinat, Nicola F. Müller, Francois-Xavier Briand, Mathieu Andraud, Axelle Scoizec, Sophie Lebouquin, Eric Niqueux, Audrey Schmitz, Beatrice Grasland, Jean-Luc Guerin, Mathilde C. Paul, Timothée Vergne

## Abstract

Phylodynamic methods have successfully been used to describe viral spread history but their applications for assessing specific control measures are rare. In 2016-17, France experienced a devastating epidemic of a highly pathogenic avian influenza virus (H5N8 clade 2.3.4.4b). Using 196 viral genomes, we conducted a phylodynamic analysis combined with generalised linear model and showed that the large-scale preventive culling of ducks significantly reduced the viral spread between *départements* (French administrative division). We also found that the virus likely spread more frequently between *départements* that shared borders, but the spread was not linked to duck transport between *départements*. Duck transport within *départements* increased the within-*département* transmission intensity, although the association was weak. Together, these results indicated that the virus spread in short-distances, either between adjacent *départements* or within *départements*. Results also suggested that the restrictions on duck transport within *départements* might not have stopped the viral spread completely. Overall, by testing specific hypothesis related to different control measures, we demonstrated that phylodynamics methods are capable of investigating the impacts of control measures on viral spread.

## Introduction

During the winter of 2016, an epidemic of the highly pathogenic avian influenza (HPAI) virus occurred in France. It resulted in 484 infected poultry farms and approximately 7 million culled birds [1]. It remains critically important to learn as much as possible from this devastating epidemic, particularly as France once again experienced an HPAI outbreak in December 2020 [2]. The 2016-17 epidemic was caused by an H5N8 virus of the lineage 2.3.4.4b (A/Gs/Gd/1/96 clade) of Asian origin that was likely introduced by migratory birds [3]. The epidemic was largely concentrated in southwest France, encompassing the Occitanie and the Nouvelle-Aquitaine *régions* (the largest French territorial administrative division) [4]. The first incidence in poultry was recorded, using clinical symptoms, in a duck breeding farm in the eastern Tarn *département* (administrative division equivalent to county or district) on 25 November 2016 and the infection was confirmed—based on lab diagnosis—on 28 November (Fig. 1). Subsequently, the epidemic spread westwards [1], following the westerly distribution of poultry farms within the region. The last case of the epidemic was reported on 23 March 2017 [1].

**Figure 1.**
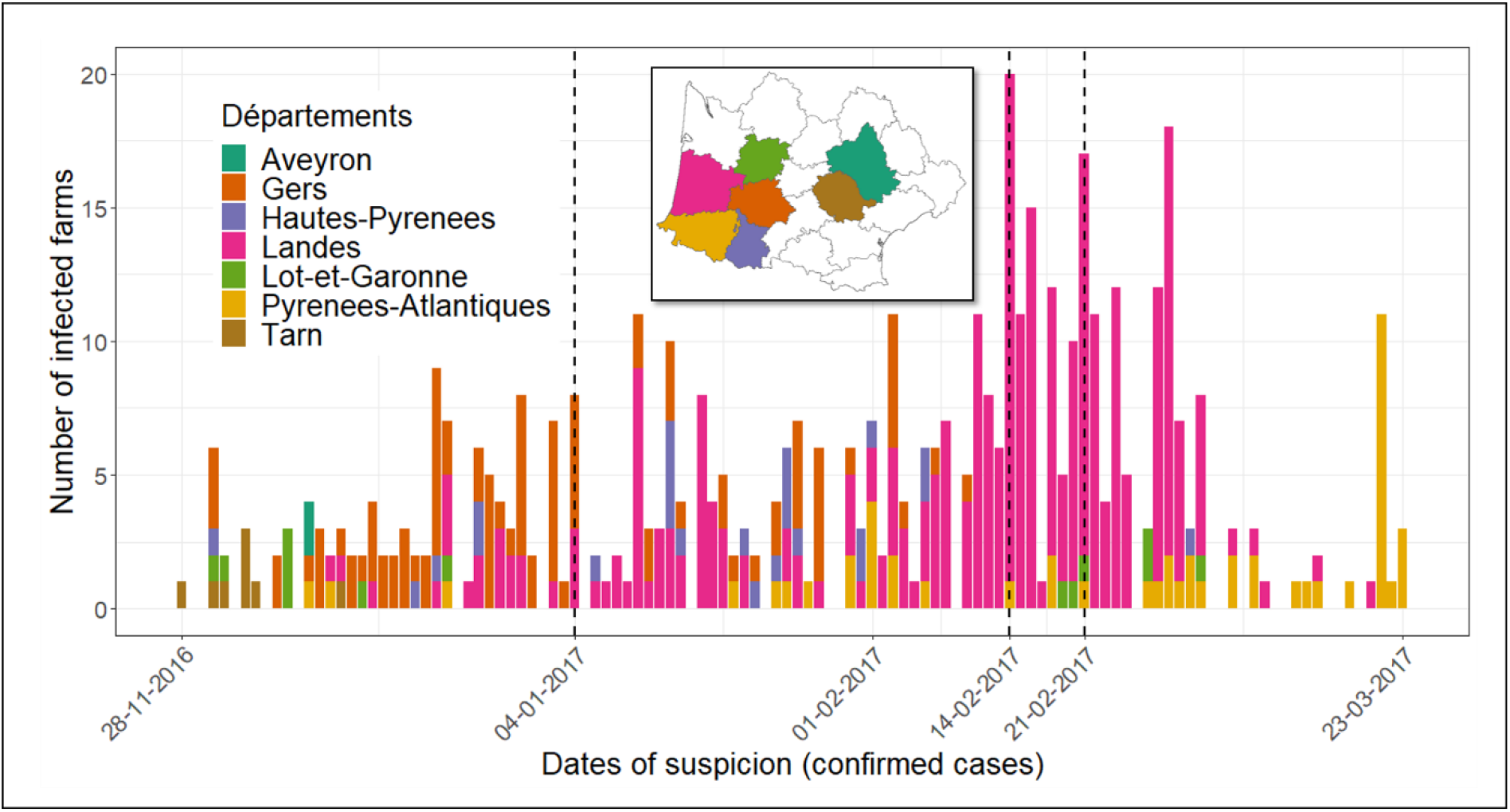
The spatio-temporal dynamics of the 2016-17 H5N8 epidemic in southwest France. The bars represent the daily number of infected farms identified in each *département*, with each *département* represented by a different colour. The number of infected farms represented those that were confirmed through PCR diagnostics during the epidemic period. The notification dates of the three culling phases are marked by three vertical lines. Inset. Location of southwest France with each study *département* marked with a specific colour corresponding to the source of samples.

Two major outbreak responses, namely preventive culling and stopping the transport of *foie gras* ducks between farms, were implemented during the epidemic. Culling was implemented locally in infected farms during the initial phase of the epidemic. However, these local measures proved to be insufficient in curbing the rapid spread of the virus. Consequently, there were three phases of large-scale culling. On 4 January 2017, all duck farms located in 150 communes across four *départements*, namely Gers, Landes, Pyrenees-Atlantiques and Hautes-Pyrenees, were notified to be preventively culled (Fig. 1). On 14 February, the number of communes under notification was doubled. On 21 February, the notified areas were increased for the last time to include more than 500 communes in total. Additionally, following detection, ducks could not be moved from or to infected farms [5].

Epidemiological studies based on incidence data have provided important insights into the HPAI epidemic patterns [1] and the control measures [5,6]. Additionally, a recent phylogenetic study analysed 196 viral sequences that were identified in poultry farms and reported that the genotype that caused the large-sale epidemic in southwest France was associated with five geographic clusters that could be the consequence of a few long-distance transmission events followed by local spread [3]. However, the ecological and epidemiological drivers of these transmission patterns are still unknown. To fill this gap, we used a phylodynamic framework that we applied to the 2016-2017 epidemic data. Viral phylodynamics is the study of viral phylogenies and how they are shaped by potential interactions between epidemiological, immunological and evolutionary processes [7,8]. These methods can detect spatio-temporal patterns of epidemics based on viral genomic data, which represent a powerful source of information and are complimentary to epidemiological data [9]. Although phylodynamics methods are frequently used for reconstructing epidemic patterns, their application in assessing control measures is still rare [10]. Here, we used a phylodynamic framework to quantify the HPAI epidemic in southwest France and tested the effect of large-scale culling, along with other potential ecological and epidemiological drivers. Doing so, we underscore the versatility of phylodynamics methods in addressing both, specific control measure-based questions, as well as more broad ones regarding viral spread history.

## Results

### Viral phylogeny and spread history

To infer the phylogeny and the geographical spread of H5N8, we used a recently developed phylodynamic model called the marginal approximation of the structured coalescent (MASCOT) [11]. It models the coalescence of lineages within and the migration of lineages between geographical units and has been applied elsewhere to study the spread of viral epidemics [12,13]. In our study, the geographical unit was the *département* as it was considered to provide a good compromise between the precision of the epidemiological inference and computational constraints, given the amount of data available. In Fig. 2A, we showed the evolutionary relationship between viral lineages from different *départements* in the form of a time-scaled summarised phylogenetic tree. Additionally, we show the most likely viral spread history of the lineages between *départements*, each of which is represented by a different colour. A change in colour across tree branches represents spread events between *départements*. We inferred Tarn to be the most likely source of all the viral lineages in southwest France (median posterior probability = 0.75, 95% credible interval; CI = 0.16-1). The median day of the emergence was estimated to be middle of November 2016 (95% CI = 27 Oct. - 22 Nov.). The clustering of basal sequences indicated a single introduction event in southwest France, followed by epidemic transmission (short terminal branches) mainly towards the west.

**Figure 2.**
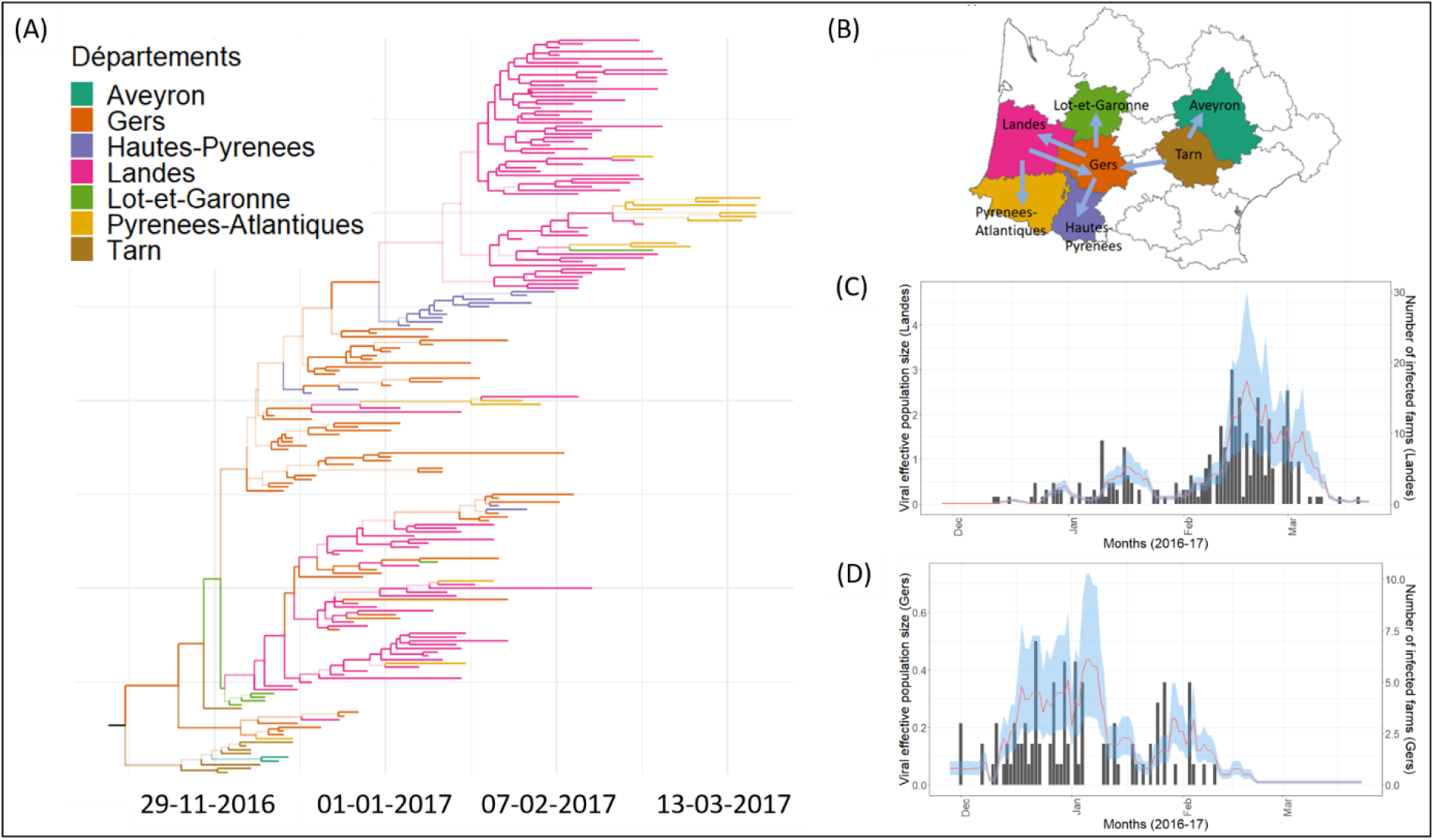
Inferred epidemics dynamics based on structured coalescent population model. (A) A time-scaled maximum clade credibility (MCC) phylogenetic tree representing the evolutionary relationship between viral lineages. The colour of a branch indicates the inferred location (see legend) with the highest posterior support. A colour change on a branch indicates a virus spread event. Numbers of all the nodes are shown in supplementary figure; (B) A schematic of the prominent (maximum probability of inferred location > 0.7) viral spread events between *départements*. (C) Estimates of viral effective population size (line in red; 95% highest posterior density; HPD in blue) and the time series of infected number of farms in Landes (black bar chart), the most affected *département*. (D) Estimates of viral population size (line in red; 95% HPD in Blue) and the time series of infected number of farms in Gers (black bar chart), the second most affected *département*.

According to the inferred migration history (Fig. 2B), the virus mostly spread between neighbouring *départements*. The only exception to this pattern was migration from Tarn to Gers. In most cases, the virus migrated in one direction, from one *département* to other and did not come back to the source. However, in a few instances—between Gers and Landes and occasionally between Landes and Pyrénées-Atlantiques, we observed bidirectional spread of the virus. Gers was the most frequent source of spread and was responsible for spread into three other *départements*, namely Lot-et-Garonne (once), Landes (in multiple occasions) and Haute-Pyrenees (once). On the other hand, both Gers and Landes were the only *départements* where viruses were introduced from more than one *département*— namely Tarn (once) and Landes (in multiple occasions) into Gers and Gers (in multiple occasions) and Pyrénées -Atlantiques (at least twice) and Landes (in multiple occasions). On the other end of the spectrum, Hautes-Pyrénées and Aveyron were the exceptional *départements* that received viruses in single events and never passed it forward to any other *départements*. Towards the end of the outbreak, lineages reached Pyrénées-Atlantiques in multiple occasions, all exclusively from Landes despite sharing its border with two other *départements* (Gers and Hautes-Pyrenees) as well.

The estimated viral effective population size *Ne* distribution for Landes—most affected in terms of case numbers—and Gers—second most affected—*départements* were proportional to the corresponding time series of case numbers (Figs. 2C and 2D).

### Identifying the predictors of viral spread between *départements*

To identify the putative predictors of viral spread between *départements*, we used the MASCOT model where the migrations rates through time are inferred with the help of a generalized linear model (GLM) and a number of potential predictors [13]. The results are shown in Fig. 3. In Fig. 3A, we show the viral spread predictors and their level of support in the form of the Bayes factor (K), which is a Bayesian alternative to classical Null-Hypothesis Significance Testing [14,15]. Predictors with at least substantial support (K > 3.2) were included in the final GLM [16]. In Fig. 3B, we show the estimated coefficients, which represents the strength and direction (positive or negative) of each predictor’s effect on the between-*département* viral spread. However, since all predictors are standardised, the strength of the effect size is not easily interpretable. The analysis demonstrated that sharing of borders (K = 3.2) as well as the period of culling of the epidemic (before and after 4 January) (K = 10) to be predictors of the observed spread patterns associated with viral spread. We inferred both predictors to positively predict spread, meaning that spread was greater between *départements* with shared borders and that spread was reduced after the 4 January culling initiation date.

**Figure 3.**
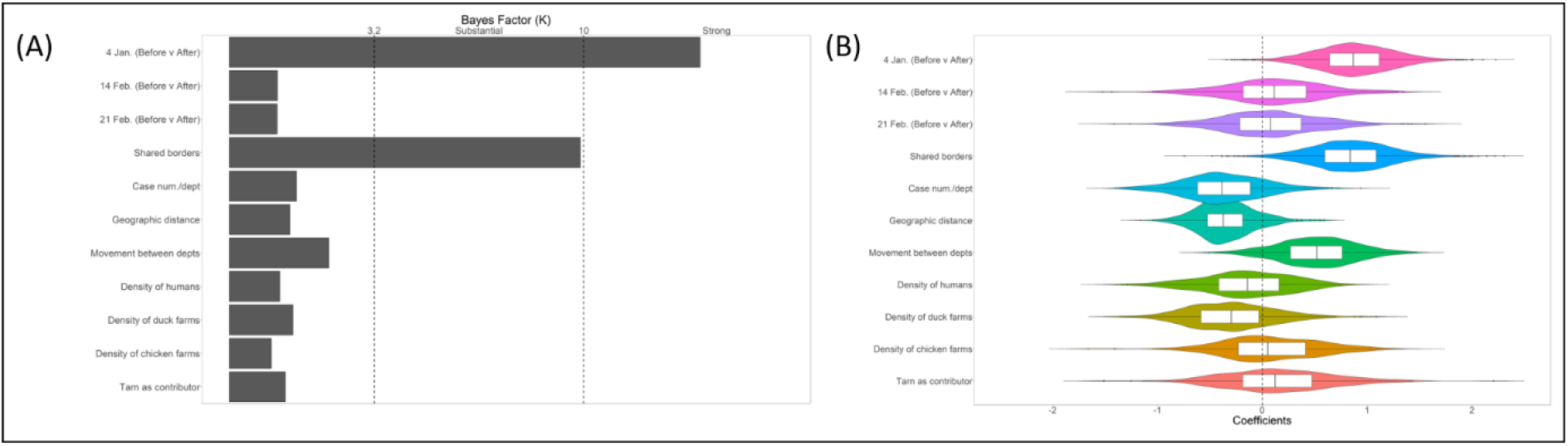
The generalised linear models (GLMs) of viral spread between *départements*. (A) The viral spread predictors and their level of support based on indicator probability, which was estimated based on Bayesian Stochastic Search Variable Selection (BSSVS). Different strengths of support are marked by vertical lines. Predictors whose indicators had at least substantial support (Bayes factor, BF = 3.2) were included in the final GLM. (B) The boxplots depict the strength and direction of each predictor’s effect.

### Identifying the predictors of viral effective population size *Ne*

To identify the putative predictors of viral effective size Ne within *départements*, we used the MASCOT model where the *Ne* through time are defined as a generalized linear model (GLM) [13]. In Fig. 4, we showed the results of a phylodynamic GLM analysis where we investigated the relationship between viral effective population size *Ne* and a number of potential predictors. In Fig. 4A, we show the *Ne* predictors and their level of support based on indicator probability. The predictors whose indicators had at least substantial support (Bayes factor, *K* = 3.2) were included in the final GLM. In Fig. 4B, we showed the estimated coefficients, which represented the strength and direction of each predictor’s effect. Based on our inclusion criterion, the final GLM again included only two predictors, namely the case numbers (K > 100, decisive) and the duck movement within *département* (K > 3.2).

**Figure 4.**
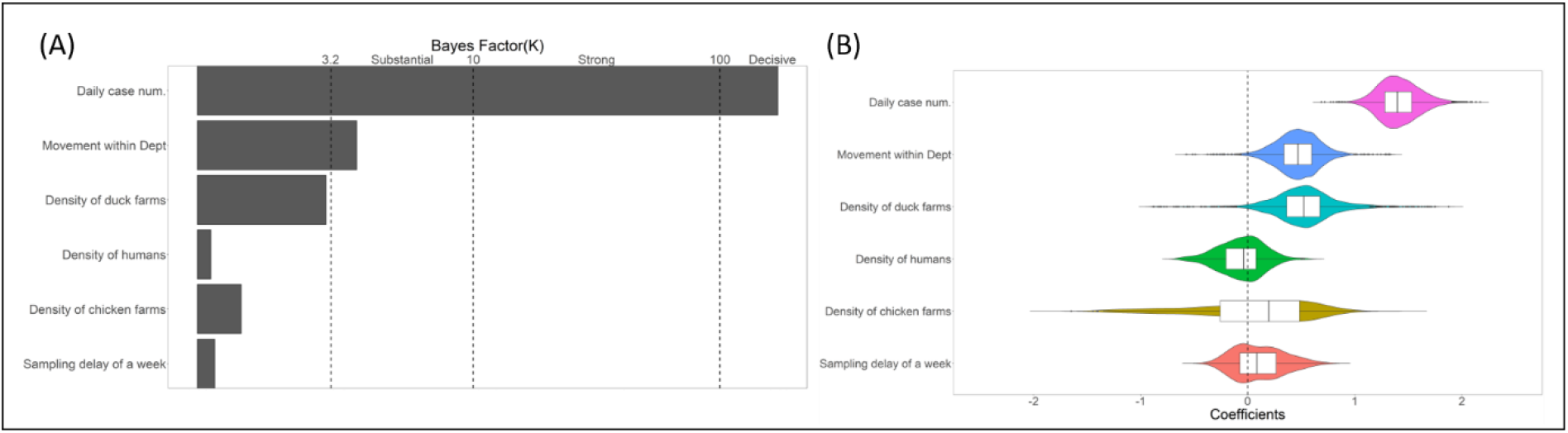
The generalised linear models (GLMs) of viral effective population size *Ne* within *départements*. (A) The *Ne* predictors and their level of support based on indicator probability, which was estimated based on Bayesian Stochastic Search Variable Selection (BSSVS). Different strengths of support are marked by vertical lines. Predictors whose indicators had at least substantial support (Bayes factor, BF = 3.2) were included in the final GLM. (B) The boxplots depict the strength and direction of each predictor’s effect.

## Discussion

In 2016-17, France experienced a devastating epidemic of highly pathogenic avian influenza (HPAI) epidemic caused by an H5N8 subtype of the lineage 2.3.4.4b (A/Gs/Gd/1/96 clade). This outbreak caused major economic losses to the industry and contributed to the implementation of stricter farm biosecurity measures [6]. Unfortunately, the virus returned to France last winter causing yet another epidemic in the south-western part of the country highlighting the importance of understanding its transmission dynamics. One of those gaps has been assessing whether control measures were effective. Employing phylodynamic methods that combine genomic and epidemiological data, we showed that the large-scale preventive culling measure initiated on 4 January 2017 was successful in reducing viral spread between *départements*. At regional scale, we showed that viral spread between *départements* occurred more frequently between *départements* that shared borders than between *départements* that were far apart, what is consistent with the geographical clusters identified in Briand et al. (2021). Our results could not find links between the viral spread between *départements* and duck transport. Within *départements*, we found that duck movements were positively but weakly associated with the viral effective population size. However, the number of infected farms was a powerful predictor of the effective population size. Additionally, our phylogeographic analysis estimated the date of viral emergence close to the date of the first detection, traced the origin of the epidemic to Tarn and exhibited detailed viral spread history between *départements*.

Preventive culling is a common control measure of viral epidemics of livestock. To reduce the epidemic size, it is often implemented in case of HPAI outbreaks because there is no treatment and vaccination is often not an option in a disease-free country like France. However, due to the elimination of potentially healthy birds and operational costs, it can be seen as an extremely expensive and controversial strategy. Therefore, it is often limited to infected farms and potentially-infected farms in the vicinity. Large-scale culling is usually implemented as a last resort to contain a rapidly spreading disease such as HPAI, after other less-expensive (e.g., local culling) and less-operationally challenging (e.g., ban on livestock movements) measures fail. Based on the HPAI epidemiologic data from the 2016-17 French outbreaks, a previous modelling study showed that a local culling strategy—culling of all poultry farms within 1 km of an infected farm–could have halved the epidemic size; but a large-scale culling strategy (up to 5 km) would only have had marginal benefits [6]. Based on this evidence and the fact that large-scale culling is operationally expensive and challenging to implement, a timely implementation of the culling of infected flocks was advocated. This finding has been echoed by other studies, both in case of HPAI [17] and foot and mouth disease (FMD) [18,19]. However, culling could also be effective in reducing viral spread, which may not be contained locally. In that respect, our results complemented the epidemiological studies by focusing on viral spread and showed that large-scale culling is useful in containing regional spread of viruses—particularly for rapidly spreading viruses such as HPAI viruses—notwithstanding its impact on the number of infected farms. Together, these results can inform culling decision-making based on the type of pathogen and epidemic stage. For instance, it would be prudent to implement local-culling strategy for a slowly-spreading virus and delay large-scale culling until absolutely necessary. But, large-scale culling could be a relevant early response for a rapidly-spreading virus as far as containing the virus is concerned. Overall, culling decision-making requires a more detailed understanding of culling strategies and their holistic impact on the virus.

In our study, we also investigated the role of the two extensions of the large-scale preventive culling, which were successively implemented on 14 and 21 February 2017. Despite including more communes for culling, our results could not find evidence that these two culling phases reduced the transmission of H5N8 beyond the effect that the first culling phase on 4 January 2017 had.

Our results did not find evidence that the amount of duck movements between *départements* impacted the spread of the virus between *départements*. This finding—along with the evidence of between-*département* viral spread only between adjacent *départements*—suggests that most live-duck movements probably did not play any role in viral spread. This is consistent with observations that the duck movements that could have been responsible for the spread of the virus were quite limited and mostly occurred at the very early phases of the epidemic before stringent movement bans were implemented [5].

This could also be explained by the implementation of a PCR-test for all duck flocks that were about to be transported. One other explanation of this pattern could be that the ban on movements successfully disrupted the association between viral spread and long-distance duck movements. However, we did not consider the duck movements as a time varying predictor, which could have impacted our ability to detect duck movements as a predictor of viral transmission between *départements*. This hypothesis could be tested in the future with duck movement data collected over time (time-dependent variable). Alternatively, our results could also mean that the virus did not spread through duck movements in the long-distance, but by some other mechanisms. Two major contenders for alternative mechanisms of viral spread are humans and farm equipment as fomites and wild birds as carriers. We did not find any significant association between human population density—which acted as a proxy for human activity—and viral spread. This suggested that human populations, which were not specifically involved in the duck-producing industry, did not play a major role in spreading the virus. An interesting development of this analysis would be to include equipment sharing data, although collection of this data is challenging as such records are probably not maintained methodically. Regarding the wild-bird related mechanisms, a study on Chinese poultry industry indicated that wild birds did not play a major role in spreading avian influenza viruses along the value chain across the country, in contrast to between-country spread for which wild birds play a prominent role [20]. This is likely to be the case in southwest France as well. Another potential mechanism could be wind-borne transmission, which is suspected in case of HPAI [21–23]. But, a previous study did not find any link between the direction of wind during the epidemic and the direction of viral spread, suggesting that the transmission was unlikely to be wind-borne [24]. However, the study was limited to a specific period during the epidemic, hence the role of wind in spreading the virus is still an open question.

Our results indicated that the virus entered southwest France through Tarn and that this event represented a single source of the epidemic. This observation is likely to be robust because i) all the samples from Tarn are monophyletic and connect to the root, ii) terminal branches are short, iii) it is consistent with a recent study based on phylogenetic approaches [3]. Although Tarn itself is not considered to have high exposure risk from migratory birds, the single introduction pattern is likely to be explained by the European routes of migratory waterfowls because the estimated emergence interval coincided with the winter migration season (October-November) of many wild waterfowl species. These species are now known to carry and spread multiple avian influenza A viruses in poultry around the world, Still, we cannot completely reject the possibility of multiple introductions of a virus on the basis of the phylogeny alone [25,26]. Indeed, if the genomic sequences from distinct introductions are similar and in turn form monophyletic clusters, then the number of introductions could be underestimated. However, given avian influenza viruses are RNA viruses with high mutations rates, this alternative hypothesis is unlikely to hold. Our finding of a single introduction that acted as the source of the whole epidemic supports a previous phylogenetic study [3] and is consistent with outbreak investigations and epidemiological analyses [1,6]. Indeed, the index case in poultry was suspected and confirmed in a duck-breeding farm in Tarn, the day after that farm sent live-ducks to force-feeding farms in Gers and Lot-et Garonne. This event is likely to have triggered the viral spread [5].

Overall, by successfully analysing the impact of control measures, we showed the versatility of viral phylodynamic methods in providing both a broad understanding of epidemics and the means to test hypotheses related to specific control measures. Finally, the methodology used in this study can be efficiently adapted to study the impact of control measures on many other viral epidemics.

## Materials and Methods

### Genomic data, bioinformatics and phylogenetic methods

196 HPAI H5N8 viruses were isolated amongst the 484 farms that experienced an outbreak between November 2016 and March 2017. The viral genomes were sequenced (one sequence per farm) at the French national reference laboratory for avian influenza (ANSES-Ploufragan, France). Each sequence was geocoded and associated with the *département* where the outbreak was located (Tarn, Aveyron, Lot-et-Garonne, Gers, Landes, Pyrénées-Atlantiques and Hautes-Pyrénées) along with the collection date. The whole concatenated virus genomic sequences were aligned using the MUSCLE multiple sequence alignment algorithm [27] implemented in MegaX software with default parameters [28]. Influenza viral genome is particularly prone to frequent genomic reassortments [29], which, if undetected, may provide spurious genomic signals. We therefore checked for the absence of recombination and reassortments in our aligned sequences using five different algorithms, namely BootScan, CHIMERA, MaxCHI, RDP and SisScan, implemented in RDP v4.1 [30]. None of the algorithms detected any recombination events. Therefore, we used all available sequences (n = 196) across the seven *départements* for downstream analyses.

We conducted an exploratory phylogenetic analysis, based on a quick maximum likelihood method, to ensure the geographic fidelity of the samples and that the samples collected earlier were closer to the root of the phylogenetic tree. This analysis justified the use of the structured and more computationally intensive models.

### Phylodynamic reconstruction of viral phylogeny and spread history

In our study, we used the GTR+Gamma_4_ substitution model which was selected as the best model by the SMS algorithm in the Programme PhyML ver. 3 [31]. We also used a strict molecular clock as is common for avian influenza viruses[32,33] and also because the maximum likelihood tree root to tip divergence and sampling dates of the samples were strongly positively correlated. The phylogenetic temporal structure was assessed using Tempest ver. 1.5.3 [34].

### Phylodynamic and generalised linear models

To infer the spread of H5N8, we used a recently developed phylodynamic model called the marginal approximation of the structured coalescent (MASCOT) [11]. This model was adapted in order to describe the coalescence of lineages within and the migration of lineages between *departments* and has been applied several times to study the spread of pathogens [12,13]. To identify putative predictors of viral spread, we used the MASCOT model where the migrations rates and effective population sizes through time are defined as a generalised linear model (GLM) [13].

### Migration rate predictors

To investigate the effects of large-scale culling of ducks, we compared viral spread before and after preventive culling was notified by French government. The assumption was that if culling was effective then viral spread should be curtailed after the date of notification. To test this scenario, we created a predictor based on 4 January 2017. This was the date of the first notification when preventive culling was implemented in 150 communes located at the border between Gers, Landes, Pyrenees-Atlantiques and Hautes-Pyrenees départements. We also created another predictor based on 14 February, when a second notification was issued to extend the culling to other communes. The last predictor of culling was based on 21 February, when the last notification was issued to include yet more communes under culling control measure. In Fig. 1, we showed the daily distribution of the number of outbreaks in each of the seven *départements* and marked the three culling notification dates.

Movement of ducks between different farms is a major feature of the *foie-gras* production in southwest France. Ducks, in large flocks of several thousands, are first raised in breeding farms for up to 12 weeks. Then, they are divided into small groups of hundreds and transported in batches to force-feeding farms across the region. Small batches are necessary because the process of force-feeding is labour-intensive and can optimally be done only in small batches. At the force-feeding-farms, the ducks are fattened for 12 days and then are sent off to slaughterhouse for harvest. As a result of this process, farms are continuously receiving and sending out shipments of ducks potentially facilitating the spread of the virus as well. To test any link between duck movement and viral spread between *départements*, we considered the total number of shipments exchanged between *départements*. This data was obtained from the professional organisation of fattening duck producers (CIFOG) for the period between October 2016 and February 2017 [35]. This predictor was considered time-independent.

Density of livestock is often linked to pathogen spread [36–38] and density of poultry farms in southwest France were found to be associated the occurrence of outbreaks in the case of the French HPAI 2016-17 epidemic [35]. To test the role of poultry density in driving viral spread, we considered the densities of duck and chicken farms per *département* as predictors. During epidemics in livestock, humans can spread pathogens between farms, either as fomites directly or through sharing of equipment [39,40]. Hence, we also considered human population per *département* as a separate predictor. These time-independent variables were computed from the French Directorate-General for Food (DGAl) and CIFOG databases [35].

We also considered the total number of infected farms in a *département* to be a potential predictor of viral spread as *département*s with high number of cases could represent a major source of virus for other departments. Also, as the first case was detected in Tarn, we considered a potential predictor that represented Tarn to the most likely source of viral spread for all *départements*.

Finally, two time-independent spatial variables were considered as previous studies identified spatial patterns in the epidemic spread [1,35]. We tested the effect of the Euclidean distance between *départements* centroids as well as the border sharing between *départements* on the viral spread between departments.

### Effective population size (Ne) predictors

In the case of Ebola virus epidemics, the number of detected cases was found to be highly predictive of the viral effective population size (*Ne*) [10,13,41]. In our analysis, we therefore used the number of infected farms in each *département* per day, which was smoothed using a 7-day moving average. However, if virus lineages are erroneously time-stamped due to unknown sampling delay, a GLM analysis may result in spurious association between case number and *Ne* [10]. So, for our second case number-based predictor, we artificially incorporated a one-week delay in the case number distribution to check for any delay [13].

To test any link between duck movement and *Ne* within a *département*, we considered the total number of shipments exchanged between farms within a *département* as another potential predictor. Livestock density plays an important role in viral transmission and hence could be linked to *Ne* [42]. We used the densities of duck and chicken farms per *département* as potential predictors. Humans, as fomites, can also influence *Ne* and hence human population density was also used as a predictor. Both these predictors were time-independent. Also, all non-binary variables were log transformed and standardised so that their means and standard deviations equalled 0 and 1, respectively.

The modelling exercises and the Markov chain Monte Carlo (MCMC) simulations were conducted in BEAST v2.6.3 [43]. For posterior sampling, we used the coupled MCMC package [44] with 100 million iterations. The convergence and mixing of the MCMC chains were checked visually in Tracer v1.7.1 [45]. An analysis was considered successful if the effective sample size of each of the parameters was at least 200. The time-scaled phylogenetic tree with the most likely *département* of each lineage was created in FigTree v1.4.3 [46].

### Selection of generalised linear model

Model selection requires extensive computation, particularly for models with many parameters. Hence, MASCOT provides an efficient model averaging approach, the Bayesian stochastic search variable selection (BSSVS), that is used to calculate a binary indicator variable [13,47]. This variable indicates if its predictor has contributed (1 = yes; 0 = No) to the GLM. BSSVS results in an estimate of the posterior inclusion probability or support for each predictor. Each predictor is also associated with a coefficient [13]. The purpose of the coefficient is to express predictor’s effect size and the value can range between -∞ to ∞. We also measured the confidence in the result based on Bayes factors (*K*). Based on convention [16], we decided the confidence to be not substantial (1 > *K* < 3.2), substantial (3.2 > *K* < 10), strong (10 > *K* < 100) or decisive (*K* > 100). In the final model, we included only the predictors with at least substantially confident result. These statistical analyses were conducted in R v4.0.3 [48]. All the data, including genetic data accession numbers, xml files and R scripts, are available in the following repository, https://github.com/dc27708/H5N8_MASCOT.

## Acknowledgements

This work was financially supported by the FEDER/*Région Occitanie Recherche et Sociétés* 2018—AI-TRACK and the PREDYT project (*Fonds pour la Recherche sur l’Influenza Aviaire*). This study was also supported by the “Chair for Avian Biosecurity”, hosted by the National Veterinary College of Toulouse and funded by the Direction Generale de l’Alimentation, Ministère de l’Agriculture et de l’Alimentation, France. NFM is funded by the Swiss National Science Foundation (P2EZP3_191891). CG is funded by the European Union’s Horizon 2020 research and innovation programme under the Marie Sklodowska-Curie grant agreement No 842621. DC gratefully acknowledges the contribution of the INRAE MaIAGE unit in the form of free access to their MIGALE computer cluster. We also thank all the members of the EPIDESA team for their support and help.

